# Evaluation of De Novo Deep Learning Models on the Protein-Sugar Interactome

**DOI:** 10.1101/2025.09.02.673778

**Authors:** Samuel W. Canner, Lei Lu, Sho S. Takeshita, Jeffrey J. Gray

## Abstract

Advances in deep learning have produced a range of models for predicting the protein-sugar interactome; however, structural docking of noncovalent protein-carbohydrate complexes remains largely unexplored. Although all-atom structure prediction models like AlphaFold3 (AF3), Boltz-1, Chai-1, DiffDock, and RosettaFold-All Atom (RFAA) were validated on protein-small molecule complexes, no benchmark or evaluation exists specifically for noncovalent protein-carbohydrate docking. To address this, we developed a high-quality dataset of experimental structures – Benchmark of CArbohydrate Protein Interactions (BCAPIN). Using BCAPIN and a novel evaluation metric, DockQC, we assessed the performance of all-atom structure prediction models on non-covalent protein-carbohydrate docking. We found all methods achieved comparable results, with an 85% success rate for structures of at least acceptable quality. However, we found that the predictive power of all models declined with increasing carbohydrate polymer length. With the capabilities and limitations assessed, we evaluated AF3’s ability to predict binding for a set of putative human carbohydrate binding and carbohydrate non-binding proteins. While current models show promise, further development is needed to enable high-confidence, high-throughput prediction of the complete protein-sugar interactome.

## Introduction

Many new computational prediction tools have recently been developed to decode the protein-sugar interactome. Bonnardel et al. created LectomeXplore, which annotates all known proteomes with a hidden Markov model (HMM) for lectins (glycan-binding proteins).^1^ If the protein is identified as a lectin, one could use Lundstrøm et al.’s model LectinOracle to predict which carbohydrate the lectin binds.^2^ However, not all carbohydrate binding proteins are lectins, for example native sugar sensors and antibodies.^3^ Leveraging this gap, some of us (Canner, Gray) developed PiCAP to predict *whether* a protein binds to carbohydrates, irrespective of protein family, and released predicted annotations on six different species, predicting a putative list of all the proteins present in the protein-sugar interactome.^4^ To further elucidate these protein-carbohydrate interactions, Canner, Shanker et al. and Bibekar et al. created CAPSIF^5^ and PeSTo-Carbs,^6^ respectively, to predict which residues a protein uses to bind to carbohydrates. The combination of these breakthrough models can be used to predict whether any given protein binds to a carbohydrate (as a lectin or non-lectin), what carbohydrate it binds to (if the protein is a lectin), and what residues are implicated in the protein for carbohydrate binding. Now, with all-atom biomolecular prediction software like AlphaFold3 (AF3),^7^ protein-carbohydrate complex structures can be readily predicted. AF3 and other deep learning models thereby make possible the development of a complete putative structural dataset of the entire protein-sugar interactome: all protein-carbohydrate interactions across a species. First however, we must evaluate the performance of AF3 and other all-atom biomolecular structure prediction models on protein-carbohydrate complexes.

The development of AlphaFold3^7^ built upon a string of advances in protein structure prediction, such as the Nobel Prize research of David Baker, Demis Hassabis, and John Jumper: Rosetta and AlphaFold2.^8^ In the past few years, the leap in the most recent generation of *de novo* prediction methods was the ability to model any molecule with all-atom structure prediction. The Jaakkola lab developed DiffDock to predict small molecule docking on a provided protein structure.^9,10^ The Baker lab developed RosettaFold-AllAtom (RFAA), becoming the first end-to-end all-atom biomolecule structure prediction.^11^ Google DeepMind released their first end-to-end all atom prediction model AlphaFold3 (AF3). Building on previous work from DiffDock, the Jaakkola lab developed Boltz-1.^12^ With partnerships from OpenAI and other industry representatives, the Chai Discovery team released their (proprietary) model Chai-1.^7,13^

Given this growing suite of models (albeit non-exhaustive), identification of their performance on specific tasks is critical, with one of the most used metrics being the success rate when benchmarked against a dataset called Posebusters.^14^ Posebusters contains non-covalent protein-small molecule complexes. Posebusters provides well-defined specificity of the small molecule and binding protein pocket, with a model’s success measured by its ability to predict small molecule complexes under 2 Å RMSD from the solved structure. In total, DiffDock and RFAA both achieve 42% success on PoseBusters,^10,11^ while AF3 and Chai-1 achieve 76% and 77% success on PoseBusters,^7,13^ respectively. No success rate was reported on PoseBusters for Boltz-1.^12^

While PoseBusters emphasizes strong specific protein-ligand binding, protein-carbohydrate interactions present unique challenges. Unlike protein-small molecule interactions, protein-carbohydrate interactions are commonly less specific, with proteins containing multiple binding sites for long linear heterogenous polymers containing various epitopes, and therefore sugars require extra attention that is not provided in the dataset.^15–17^ Moreover, proteins stabilize carbohydrates through a combination of direct contacts (hydrogen bonding, electrostatics), indirect (water mediated) interactions, and by CH-π bonds via aromatic residues.^17,18^ Finally, the binding affinity of protein-carbohydrate complexes are commonly weak (μM – mM), but rather driven by high avidity (nM) of multiple binding sites on the protein or multiple repeats of the glycan epitope.^3^

Due to the distinct binding mechanisms involved in noncovalent protein-carbohydrate interactions, solved experimental structures of bound non-covalent carbohydrates to proteins are limited. From all solved structures in the Protein Data Bank, DIONYSUS identifies protein structures with non-covalent specific interactions with carbohydrates to be 2.5% (5,461).^19,20^ With the advent of high-throughput diazirine photoaffinity linker experimental data of protein-carbohydrate interactions,^21,22^ we are attaining more knowledge of protein-carbohydrate interactions on a protein level. We therefore propose that all-atom deep learning (DL) structure prediction pipeline may enhance our understanding of the protein-sugar interactome (PSI).

Here, we benchmark DL structural models: AF3, Boltz-1, Chai-1, RFAA, and DiffDock on the task of predicting docked *de novo* protein-carbohydrate structures. To benchmark the models, we constructed a novel dataset of proteins unseen during each model’s training. We identify the strengths and shortcomings of these models and evaluate test cases where all models perform poorly. With strengths and limitations identified, we then use AF3 as a proof-of-concept tool for predicting the structural *de novo* human protein-sugar interactome. This work sets the stage for future integration of deep learning tools in structural glycobiology to fully characterize the protein-sugar interactome across all species.

## Results

### BCAPIN and DockQC: Novel datasets and analysis

To assess the capabilities of AlphaFold3 (AF3), Boltz-1, Chai-1, DiffDock, and RosettaFold All-Atom (RFAA) at predicting protein-carbohydrate complexes, it is essential to have an independent test set of high-quality experimentally resolved protein-carbohydrate structures and a suitable evaluation metric. For the dataset, we leveraged DIONYSUS,^20^ which aggregates all experimentally determined protein-glycan structures from the PDB. We first excluded all protein-nucleic acid complexes and clustered the remaining protein sequences at 50% identity. We removed clusters with structures solved before the latest model’s training cutoff dataset (September 2021). Importantly, due to experimental limitations, not all experimental structures are of equal quality. To ensure structural reliability, we applied a filter using the real space correlation coefficient (RSCC)^23^, which measures the agreement between the calculated and experimental density. Structures with an RSCC greater than or equal to 0.9 were retained (Figure S2). The resulting Benchmark of CArbohydrate Protein INteractions (BCAPIN) test set consists of 20 structures: 9 structures that bind sugar monomers, 3 structures that bind dimers, 5 structures that bind polymers, and 3 structures that bind at least a nucleotide (NTP) and a saccharide (Table 1).

**Table 1:**
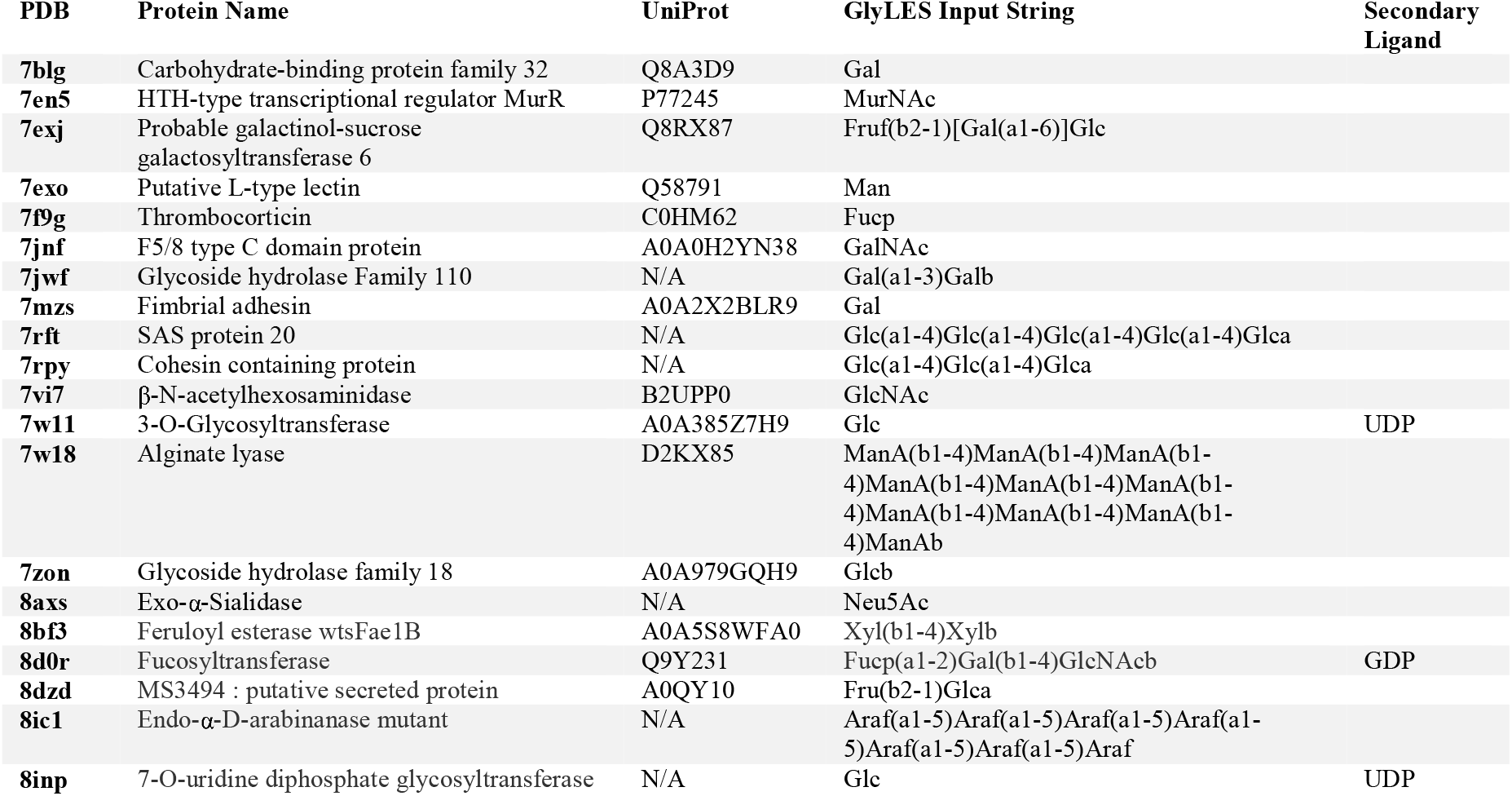
Benchmark of CArbohydrate Protein INteractions (BCAPIN) test set. The table lists the PDB 4-letter ID, protein name, UniProt ID, glycan input string for GlyLES, and any secondary ligands if present.

To evaluate the performance of predicted protein-carbohydrate complexes, we developed a single continuous scoring metric named DockQC. DockQC is inspired by the DockQ metric from the CASP-CAPRI challenge, averaging the fraction of native contacts (*F*_nat_), interface root mean squared deviation (IRMS), and ligand RMS (LRMS) to designate a predicted structure’s quality. While DockQ is widely used for protein-protein docking, the native code is unusable on our test cases, and, when reimplemented, it tends to overestimate the quality for protein-carbohydrate complexes, often assigning medium-to-high scores even when the predicted ligand position is incorrect (Figure S1, Table S1).

DockQC addresses these issues by averaging three terms: *F*_nat_, ring-ring RMSD (rRMS), and LRMS. *F*_*nat*_ measures the fraction of native residue-residue contacts, *rRMS* is a novel metric that measures the RMSD between the center of mass (COM) of each carbohydrate ring in the aligned predicted and experimental structures, and *LRMS* measures the RMSD of all aligned ligand heavy atoms.

With the BCAPIN test set and evaluation metrics established, we investigated the performance of five methods, AlphaFold3, (AF3), Boltz-1, Chai-1, RosettaFold All-Atom (RFAA), and DiffDock, at predicting protein-carbohydrate structure. We first evaluated the behavior of DockQC on the set. Thresholds were chosen after inspecting many predictions and tuning metric weights, some examples are described next.

On hedgehog interacting protein (7PGK), which binds a disaccharide heparin analog, Chai-1 failed to predict the protein structure accurately, leading to an incorrect carbohydrate placement with a low DockQC score of 0.11 (Figure 1A). For chitoporin (7EQR), a β-barrel protein that binds an oligosaccharide with a degree of polymerization (DP) of 1 six, RFAA captured the binding pocket of the carbohydrate, but lacked broader structural accuracy, yielding an acceptable prediction a DockQC of 0.26 (Figure 1B). With sialidase-sialic acid complex (8AXS), Boltz-1 achieved a medium quality prediction, correctly modeling the binding pocket and ring position (but not its orientation), with a DockQC of 0.65 (Figure 1C). In contrast, on glycoside hydrolase family 110 protein binding a Gal dimer (7JWF), AF3 nearly recapitulated the experimental structure delivering a high-quality structure with a 0.96 DockQC (Figure 1D). In total, our DockQC quality thresholds chosen to be incorrect (DockQC < 0.25), acceptable (0.25 <= DockQC < 0.50), medium (0.50 <= DockQC < 0.80), and high (DockQC >= 0.80) (Figure S1, Table S1).

**Figure 1:**
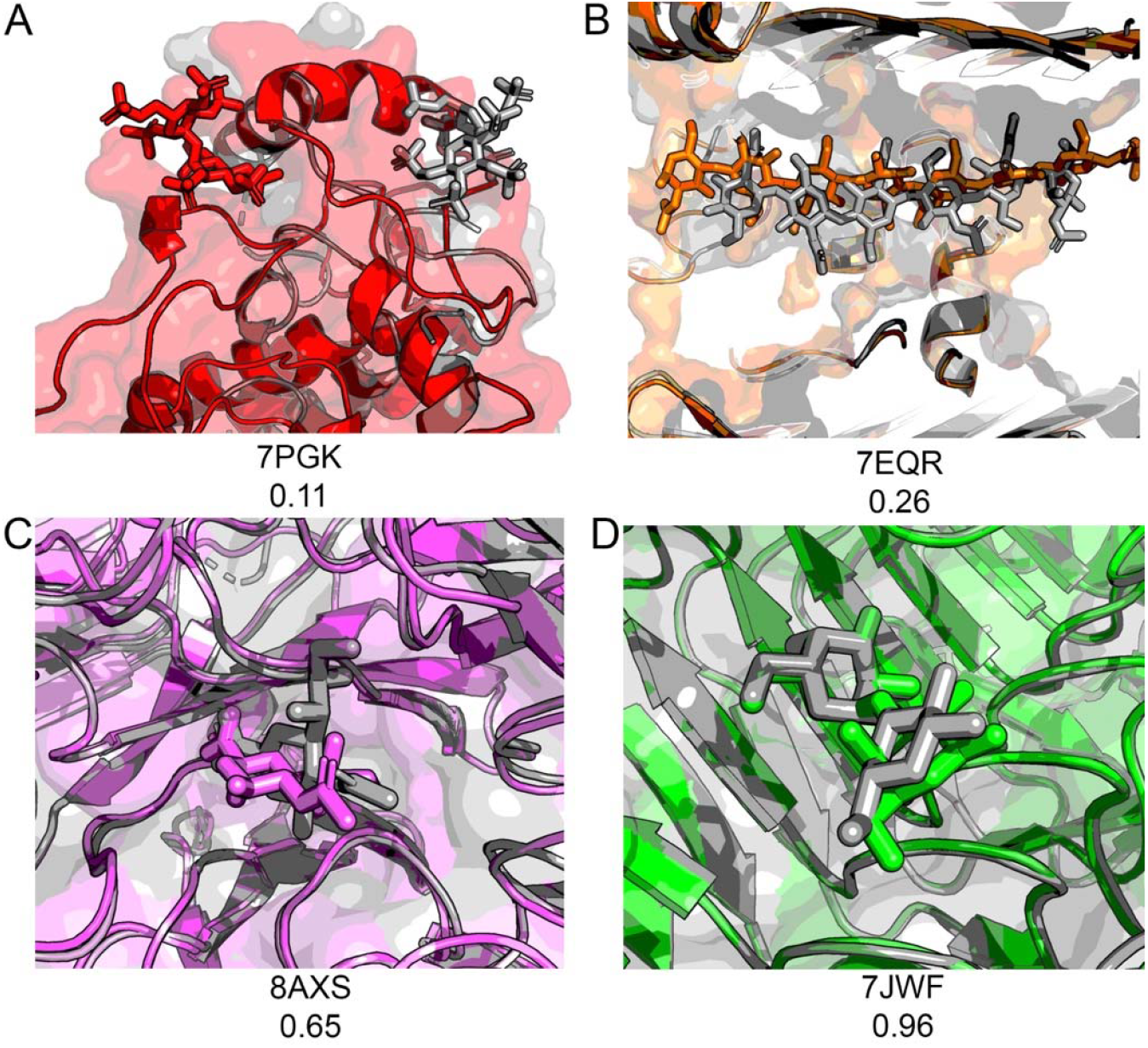
Protein-carbohydrate docked structures across DL methods. (A) Incorrect prediction of Chai-1 (red) on 7PGK (DockQC = 0.11). (B) Acceptable quality prediction of RFAA (orange) on 7EQR (DockQC = 0.26). (C) Medium quality prediction of Boltz-1 (violet) on 8AXS (DockQC=0.65). (D) High quality prediction of AF3 (green) on 7JWF (DockQC=0.96)

### DL Methods achieve medium or high accuracy on over 80% of cases

After tuning our DockQC metric, we evaluated overall model performances on all BCAPIN targets (Figure 2). Across methods, we found comparable results for all end-to-end models, at least 80% of their the highest confidence predictions (top-1) scored with at least acceptable quality. Expanding scoring to include the most accurate of each model’s top 5 confidence predictions (top-5) led to only marginal improvements. AF3 was the best-performing model: its top-1 predictions yielded 10% acceptable, 40% medium, and 35% high quality structures; top-5 predictions improved slightly to 15% acceptable, 35% medium, and 40% high quality structures.

**Figure 2:**
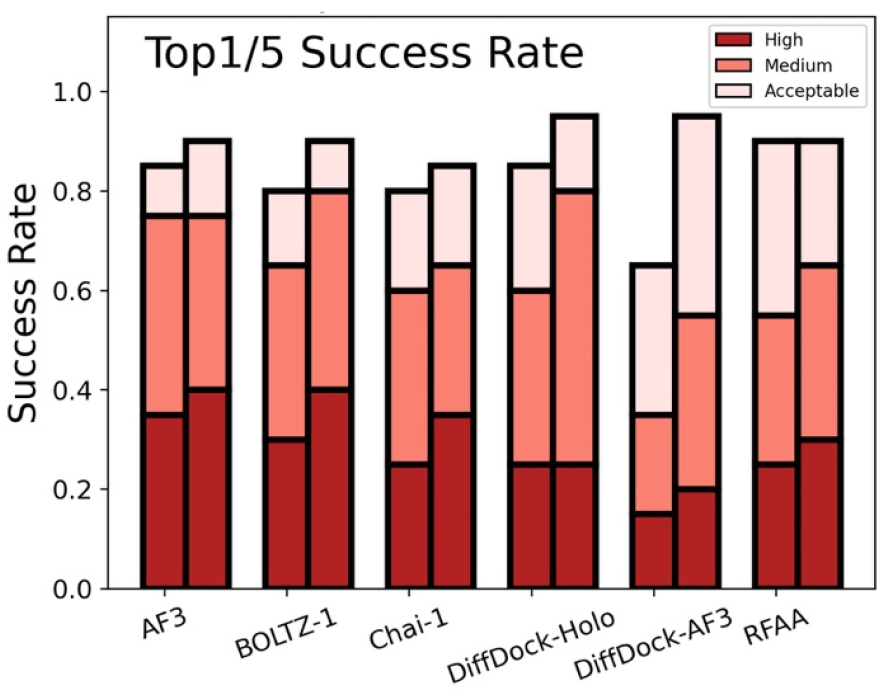
DL model success rates on BCAPIN Test Set. Each labeled method has the top-1 model on the left and top-5 model on the right.

Given the strong performance of end-to-end models on BCAPIN, we next examined how starting structure influences DiffDock’s predictive power. DiffDock-*holo* (initialized with the experimentally solved *holo* protein structure) performed equivalently to the end-to-end models, achieving at least acceptable quality on 85% of all top-1 predictions. In contrast, Diffdock-*AF3* (initialized from AF3-predicted *apo* protein) achieved only 60% acceptable or better quality in top-1 predictions. However, when extending to the top-5 predictions, Diffdock-AF3 improved substantially, yielding 85% acceptable quality structures. Thus, DiffDock is sensitive to the initial input structure.

### Methods fail to capture all cases

Although all models perform strongly on BCAPIN, we sought to identify cases where all models still struggle. Notably, all models fail to predict on two complexes: 8DZD and 7ZON (Figure 3).

**Figure 3:**
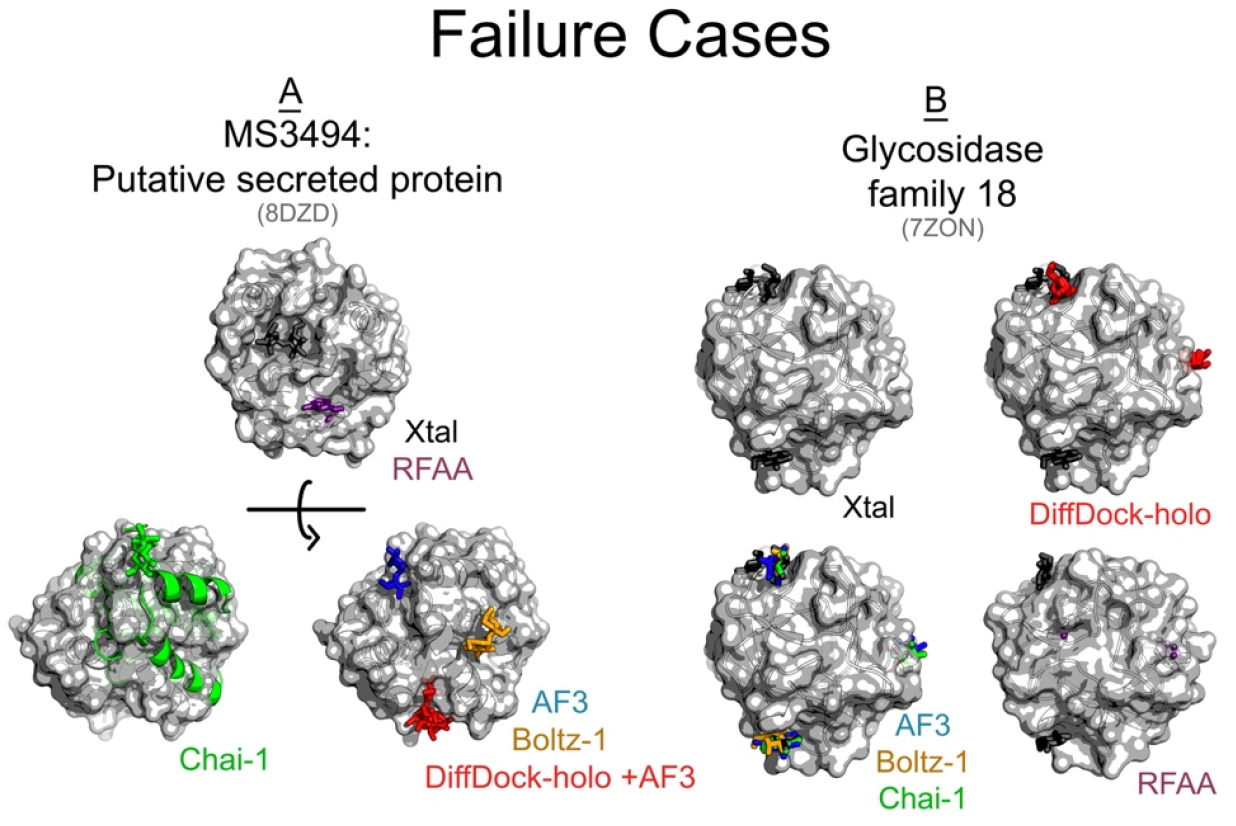
Failure of DL prediction algorithms on select proteins from the BCAPIN test set. Experimentally solved structures of (A) secreted protein (8DZD) and (B) glycosidase family 18 (7ZON, right) in gray, alongside AF3 predictions (blue), Boltz-1 (orange), Chai-1 (green), Diffdock (red), and RFAA (magenta).

8DZD is a *Mycobacterium smegmatis* secreted protein composed entirely of α-helices bound to a fructose-glucose disaccharide. While most models (except Chai-1) accurately predict the protein backbone, none correctly dock the ligand. RFAA places the ligand inside the protein. 7ZON is a glycosidase primarily composed of β-sheets bound to three independent glucose monosaccharides. Although most models correctly predict two of the binding sites, the models consistently misplace the third monosaccharide on the opposite side of the protein surface.

We further scrutinized all predictions to identify additional cases of sub-optimal performance. We found that all models produced only acceptable to medium quality on 8IC1 and 7RFT (Figure 4).

**Figure 4:**
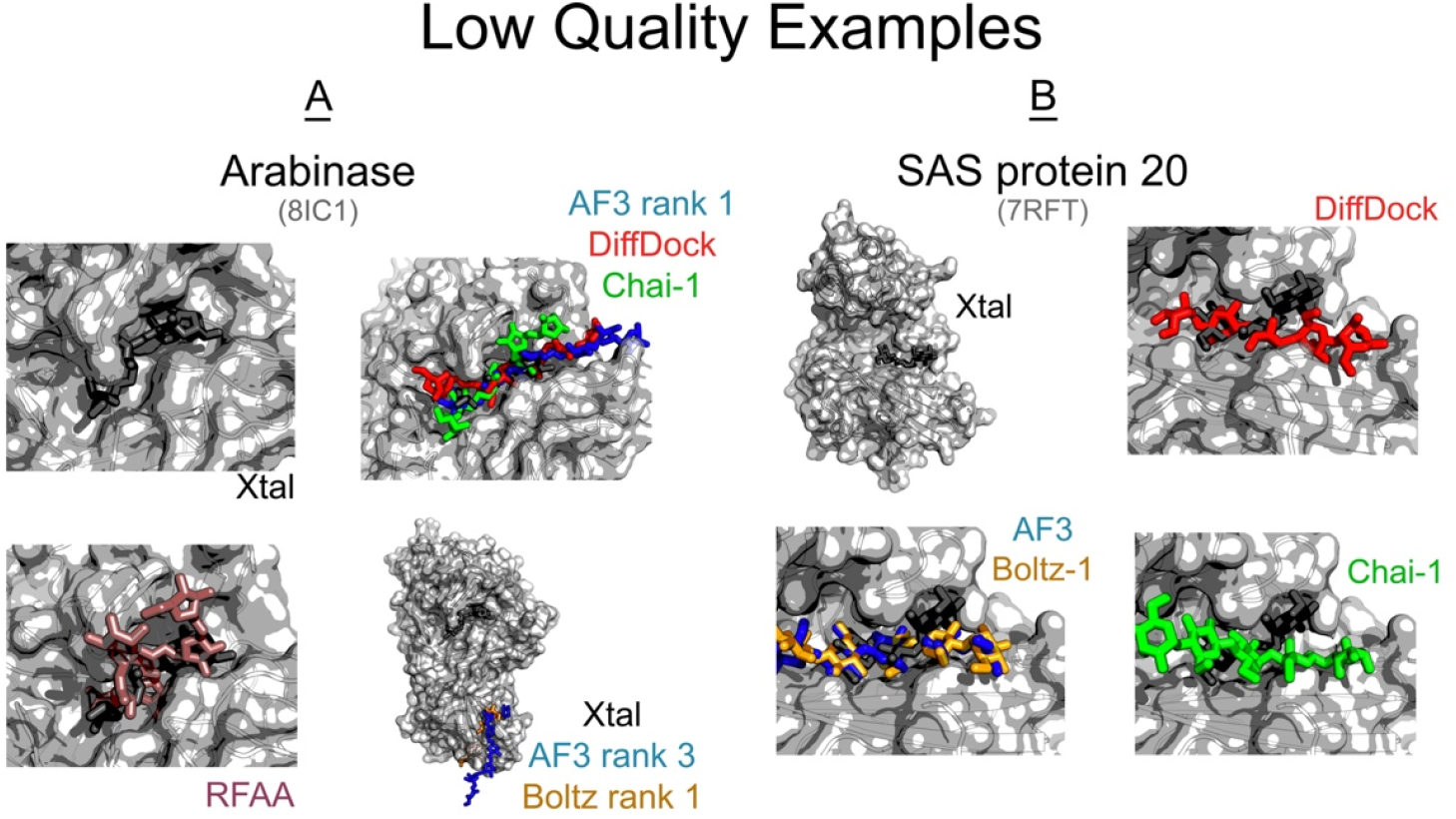
Low Quality DL predictions select proteins from the BCAPIN test set. We show the experimentally solved structures of (A) arabinose (8IC1) and (B) SAS protein 20 (7RFT), in gray, alongside AF3 predictions (blue), Boltz-1 (orange), Chai-1 (green), Diffdock (red), and RFAA (magenta).

8IC1 is an arabinose that binds a homogenous arabinofuranose oligosaccharide of DP 4 along a β-sheet. Several models, such as AF3 and Boltz-1, incorrectly predict binding at an alternative β-sheet, while others (DiffDock and RFAA) incorrectly predict the saccharide conformation (Figure 4A). 7RFT is a SAS protein 20 that binds a glucose oligosaccharide of DP 3 at a β-sheet. Although all methods identify the binding pocket of 7RFT correctly, none accurately reproduce the specific experimental conformation, particularly the orientation of the terminal Glc, which experimentally makes minimal contact with the β-strand (Figure 4B). These data suggest that current models may have difficulty on α-helical binding pockets of saccharides, simultaneous binding of multiple ligands, and docking longer saccharides.

### Prediction power decreases with carbohydrate length

We next hypothesized that model performance may correlate with saccharide complexity. To explore the role of DP on performance, we plotted the top-1 DockQC score against saccharide length (Figure 5). In total, all models showed similar trends across saccharide length categories: medium quality for monosaccharides, medium to high quality for disaccharides, acceptable quality for oligosaccharides, and acceptable quality for glycosyltransferases (GTs). Thus, we observed a decline in performance as complexity increased from simple mono and disaccharides to DP of three or greater and coordination of small ligands, in the case of GTs.

**Figure 5:**
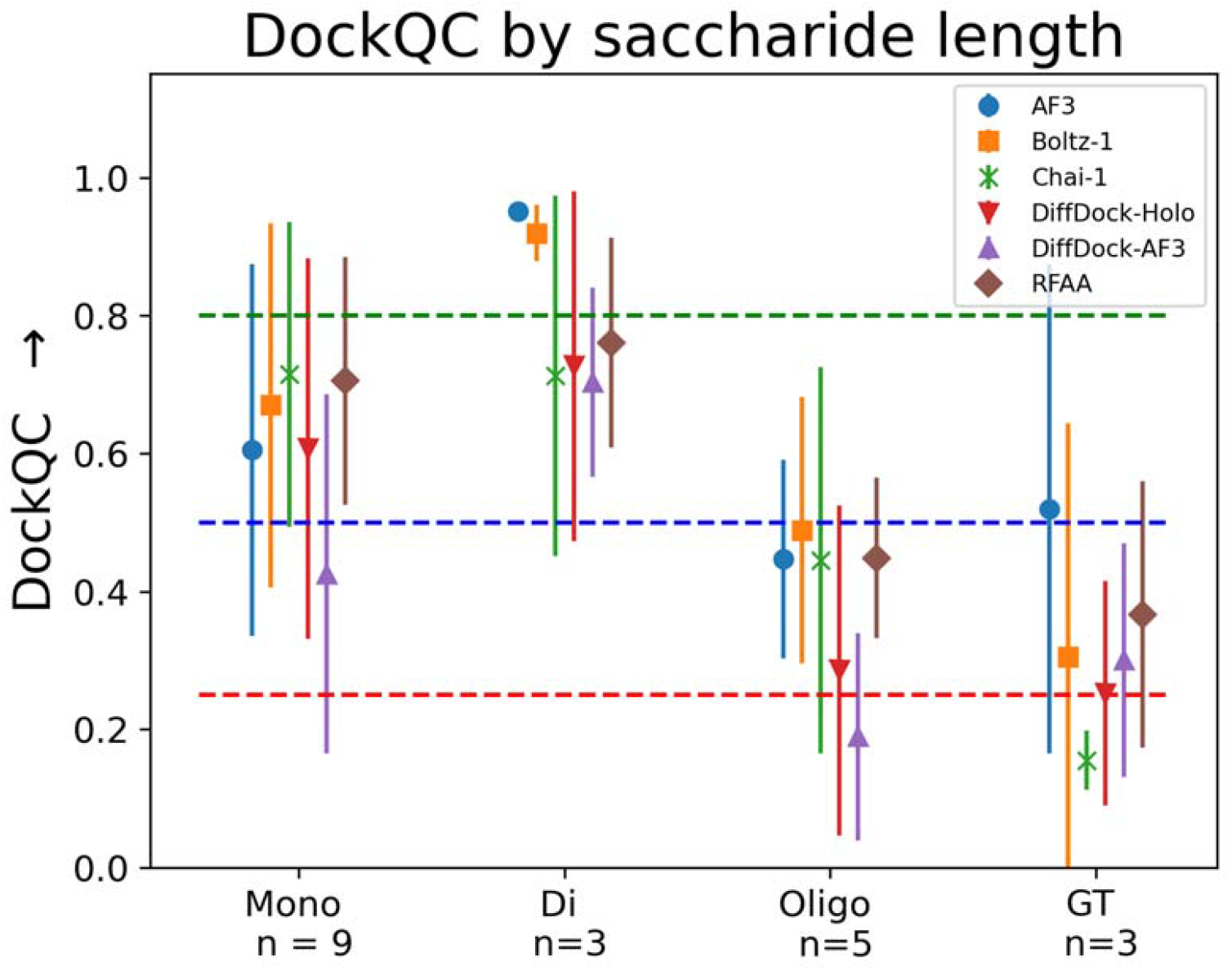
Comparison of average and standard deviation DockQC of predicted structures versus saccharide length. We group saccharide length into a degree of polymerization (DP) of 1 (mono), 2 (di), and 3+ (oligo), and further group all glycosyltransferases (GTs) together that require multiple inputs (e.g. a saccharide and NTP) and with the number of proteins in each group listed. Dashed lines indicate the DockQC cutoffs between acceptable (red), medium (blue), and high (green) quality structures. Top-1 prediction on BCAPIN with AF3 (blue circle), Boltz-1 (orange square), Chai-1 (Green X), Diffdock-*holo* (red triangle), Diffdock-*AF3* (purple triangle), and RFAA (brown diamond).

**Figure 6:**
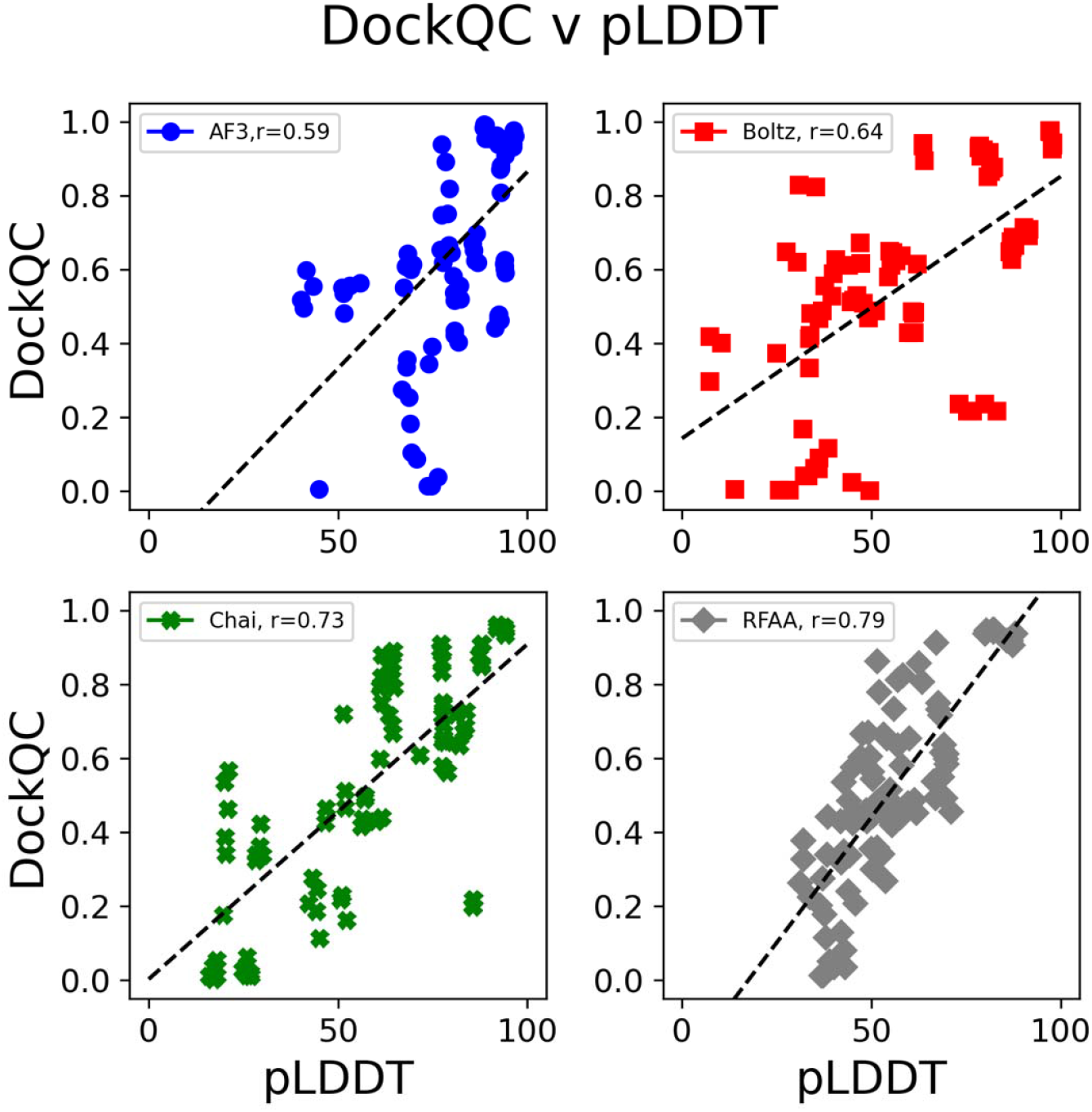
Comparison of confidence metrics and DockQC accuracy on the BCAPIN test set. Lines of best fit are provided for each plot. (A) Comparison of DockQC and ligand pLDDT for AF3 (blue circle), Boltz-1 (red square), Chai-1 (green X) and RFAA (gray diamond). (B) Comparison of DiffDock confidence for both DiffDock-*holo* (red circle) and DiffDock-*AF3* (blue square). (C) Comparison of DockQC and ipTM for AF3, Boltz-1, and Chai-1. (D) Comparison of DockQC versus the pAE for AF3, Boltz-1 (called pDE), and RFAA.

### Model confidence moderately predicts accuracy

Although all current models perform strongly on BCAPIN, performance varies across predictions. We therefore assessed whether models can reliably self-assess the accuracy of their own predictions using internal confidence metrics, such as predicted local distance difference test (pLDDT), interface predicted template modeling score (ipTM), and predicted absolute error (PAE). For average ligand pLDDT, AF3 and Boltz-1 show moderate correlations with DockQC, whereas Chai-1 and RFAA produce strong correlations (Figure 4). Since pLDDT reflects only the ligand confidence, we also evaluated ipTM, which incorporates the protein-ligand interface. Among models reporting ipTM (AF3, Boltz-1, Chai-1), all show moderate correlations, with Chai-1 performing best (Supplemental Figure S3). For PAE, Boltz-1 showed a weak negative of -0.26, AF3 a moderate correlation, with RFAA a strong correlation of -0.7 with DockQC (Supplemental Figure S4).

Contrary to the end-to-end models, DiffDock provides only one confidence metric. While both DiffDock-holo and DiffDock-AF3 use the same scoring, DiffDock-*AF3*’s provides a significantly weaker correlation than DiffDock-*holo*, reinforcing DiffDock’s sensitivity to the starting structure (Supplemental Figure S5).

Overall, all end-to-end models show moderate correlations between their internal confidence metrics to the DockQC, with RFAA demonstrating the strongest predictive reliability. Contrarily, DiffDock’s confidence metric is more susceptible to small perturbations in the input structure, limiting its reliability.

### Proteome scale predictions require refinement

The BCAPIN dataset is limited to small (less than 600 residues) single- or two-domain structurally resolved proteins with strong binding affinities. Despite being implicated in important physiological interactions, binding characteristics of large multidomain or multichain structures with carbohydrates are less well characterized due to their relative low binding affinity (but high avidity). To elucidate the protein-sugar interactome, researchers currently employ photoaffinity tag experiments^22^ or use computational tools like LectinOracle^2^ or PiCAP.^4^ However, these tools do not provide structural protein-carbohydrate complex predictions. Therefore, we aimed to assess if any end-to-end all atom structure prediction models could provide a high-throughput *de novo* approach for predicting docked protein-carbohydrate complexes with high confidence. To evaluate a *de novo* protein-carbohydrate docking pipeline, we selected nine proteins from the human proteome and used AF3 with its ipTM confidence metric to predict their structures in complex with either a GM1 ganglioside or a hybrid N-glycan (Figure 7). We used GM1 ganglioside ligands for proteins experimentally identified to interact with GM1 gangliosides in Zhang et al. and the hybrid N-glycan ligand for all others, as it is a common covalent modification on membrane and secreted proteins.

**Figure 7:**
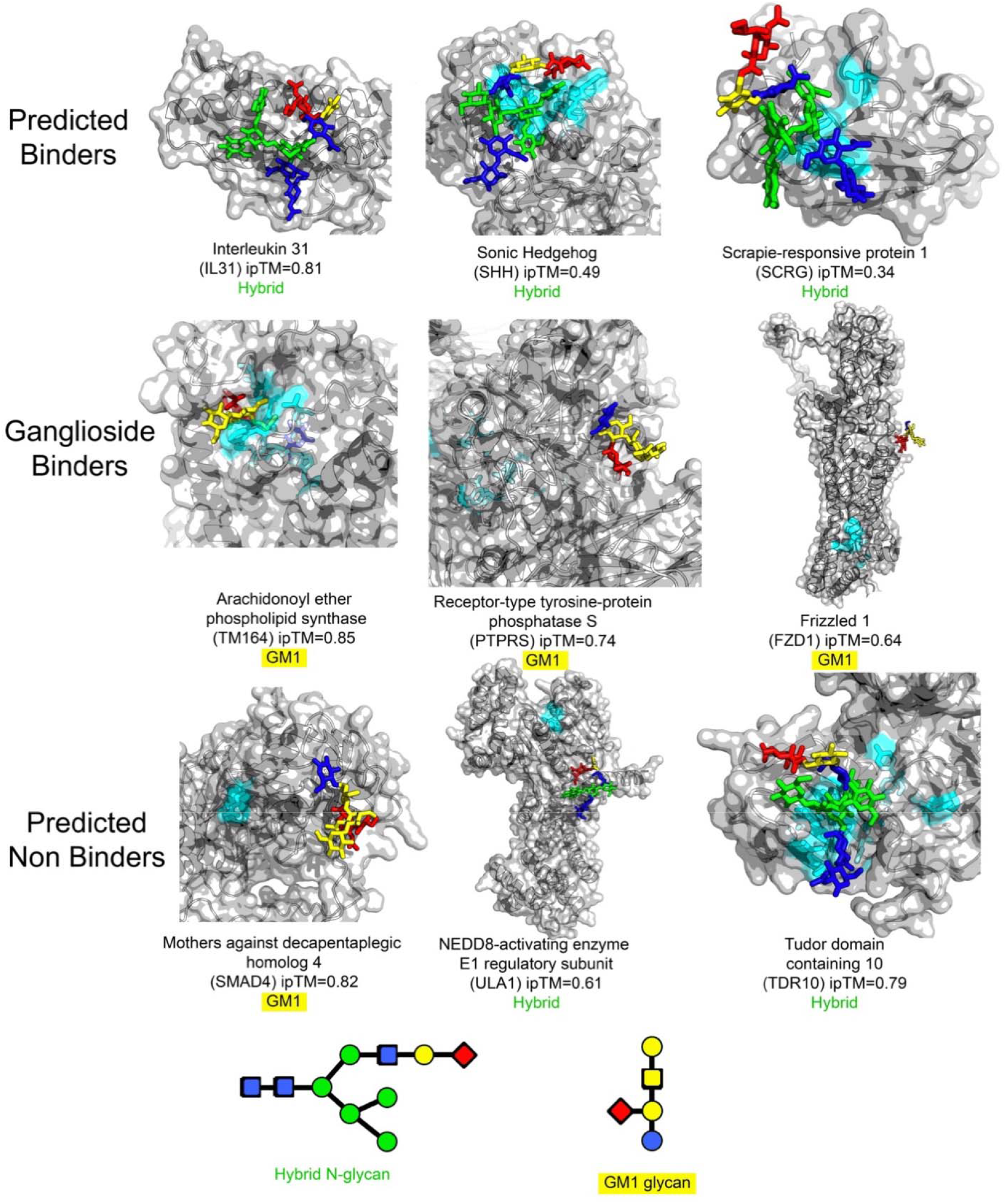
AlphaFold3 predictions on selected human protein-glycan interactions. PiCAP provides the protein-level prediction, and CAPSIF2 provides residue predictions (cyan). The bound glycan is either a complex N-glycan (green) or a GM1 ganglioside (yellow), with the initial GlcNAc of the N-glycan highlighted in blue and all sialic acids highlighted in magenta.

PiCAP predicted interleukin 31 (IL31), sonic hedgehog (SHH), and scrapie-responsive protein 1 (SCRG) as putative carbohydrate binding proteins. Here, we used AF3 to dock these proteins with a hybrid N-glycan, a common branched saccharide where one branch terminates in an oligomannose chain and the other in a sialic acid. CAPSIF2 predicted no carbohydrate-binding residues on IL31; however, AF3 predicted the glycan to bind at an unstructured region of the protein with a high interaction confidence (ipTM = 0.81). Conversely, AF3 docked the N-glycan at the CAPSIF2 predicted residues of SHH and SCRG with a lower confidence (ipTM = 0.49).

Experimentally, arachindonyl ether phospholipid synthase (TM164), receptor-type tyrosine-protein phosphatase S (PTPRS), and Frizzled 1 (FZD1) were identified in multiple experiments as ganglioside binding proteins.^22^ These proteins were also predicted by PiCAP to bind a carbohydrate. We therefore modeled these proteins in complex with the GM1 ganglioside glycan (Figure 7). AF3 predicted TM164 to bind GM1 in the CAPSIF2 predicted pocket with high confidence (ipTM = 0.85). However, AF3 however predicts PTPRS and FZD1 to bind the ganglioside glycan at sites outside of the CAPSIF2 predicted pockets. Notably, CAPSIF2 predicts on intracellular binding pocket for FZD1, whereas both AF3, experimental data, and CAPSIF:V suggest binding occurs in the extracellular region.^22^

While PiCAP predicts approximately 7,000 human proteins to bind carbohydrates, it also predicts ∼13,000 human proteins as non-binders. To assess whether AF3 could also discriminate between physiologically relevant and irrelevant interactions, we selected three proteins: mothers against decapentaplegic homolog 4 (SMAD4), NEDD-8 activating enzyme E1 regulatory subunit (ULA1), and Tudor domain containing 10 (TDR10). Since SMAD4 was previously investigated by Zhang et al. and identified as a putative *non*-binder of GM1, we modeled the protein with GM1. AF3 however predicts the SMAD4-GM1 complex with a high confidence (ipTM = 0.82). Similarly, AF3 predicted moderate to high confidence interactions for an N-glycan in complex with ULA1 (ipTM = 0.61) and TDR10 (ipTM = 0.79). These findings suggest that ipTM values alone may not be sufficient to distinguish between physiologic and non-physiologic interactions in a high-throughput manner.

## Discussion

We present an evaluation of multiple end-to-end all-atom prediction frameworks for carbohydrate-protein docking and interrogate their capabilities at unveiling the structural secrets of the protein-sugar interactome. Overall, all methods perform incredibly well at this task – all end-to-end models capture 80% of their highest confidence models at least acceptable quality (Figure 2). These models improve upon previous energy-based protein-carbohydrate docking methods like GlycanDock^24^ and HADDOCK^25^, which are useful for refinement but not full *de novo* docking. Although the models we tested improve upon previous methods and models, they still have limitations, including reduced performance with increased complexity. Specifically, the models perform worse on multi-ligand targets (GTs) and saccharides with DP greater or equal to three. Also, the models lack robust confidence metrics for protein-carbohydrate complexes.

Our BCAPIN dataset is the first study of protein-carbohydrate noncovalent docking, including all protein-carbohydrate complexes in the PDB. However, BCAPIN primarily comprises small, globular, single-domain proteins bound to linear glycan chains, which is not representative of the diverse protein-carbohydrate interactions found in physiological contexts. Thus, as more experimental data becomes available, alongside further developments in these prediction techniques, the framework presented here can be iterated to better elucidate the protein-sugar interactome.

Several limitations are present in our work. Many protein-oligosaccharide complexes are dominated by non-specific electrostatic interactions. In some instances, a few central residues engage in specific interactions while terminal residues interact primarily through electrostatics, resulting in reduced experimental resolution, as seen in 7PUG and 7EQR in our low quality test set (Table S2). Other cases, like heparin-binding proteins, are driven almost exclusively by electrostatic forces.^26^ To address these issues, we applied an average RSCC cutoff on bound ligands, though this approach removes several high-quality specific bound structures with unstructured termini. Additionally, BCAPIN does not include heparin binding proteins, which should be studied for future work.

The largest limitation in continually iterating and benchmarking this structure prediction software however is the availability of high-quality experimental structures. Although the DIONYSUS dataset is impressive in its scope, containing 5,461 protein-carbohydrate complexes, only 1,842 unique protein structures remain after by 95% sequence similarity^20,27^. Further, when assessing the individual unique binding pockets of these DIONYSUS proteins, there are only 258 unique clusters of binding pockets.^28^ With this limited set of ∼1,800 unique structures and ∼250 unique binding mechanisms, data science and machine learning approaches are restricted. Therefore, discovery of novel carbohydrate binding proteins and their structural interactions is critical.

To better improve computational approaches, we believe that one of the most promising sources of future data future lies in liquid glycan arrays and photoaffinity labeling experiments (e.g. those using diazirine linkers).^21,22,29,30^ These *in vivo* high throughput techniques enable identification of protein-carbohydrate interactions on a proteome-wide scale; however, they currently lack immediate structural resolution. Computational modeling stands poised to fill this gap by providing structural hypotheses at atomic level detail, thereby accelerating the validation and functional understanding of these experimentally identified interactions. To push the scope of the BCAPIN test set, we selected two branched polysaccharides with distinct properties to explore AF3’s capabilities. Although our study does not demonstrate that AF3 is yet ready to support full scale high-throughput experiments comparable to photoaffinity labeling, it shows that AF3 can generate useful, testable hypotheses on a case-by-case basis that may expedite wet lab investigations.

To aid wet lab experiments, our lab has computationally studied protein-carbohydrate structural interactions. We developed GlycanDock^24^, CAPSIF^5^, and PiCAP^4^ as ways to elucidate these interactions. PiCAP in particular, represents a significant advancement, as it was the first model to predict *whether* a protein binds to carbohydrate, irrespective of protein family on a proteome scale. However, these current models rely on the fundamental work of thousands of scientists solving crystal structures of protein-carbohydrate complexes. While high-throughput technologies are likely to uncover many more non-covalent protein carbohydrate interactions *in vivo*, reliably obtaining the bound structure or identifying the full glycan repertoire for each protein remains a computational bottleneck.

We envision a full suite of models and methods will fill the gap to identify the full protein sugar interactome of a species. We advocate for a model that would improve upon LectinOracle^2^, integrating the glycan embeddings from methods like SweetNet^31^ or Gifflar^32^ using sequence and structural information insights from structure prediction models, current photoaffinity experiments, and CAZY^33^ can predict the glycan binding repertoire of all proteins. With this addition, one can use PiCAP to predict whether a protein binds carbohydrates, use CAPSIF2 or PeSTo-Carbs to predict how the protein binds the carbohydrate structurally, and finally, use the proposed model to predict which carbohydrates are recognized, all at high-throughput scales. This integrated approach will be essential to fully map the protein-sugar interactome, advancing our understanding of glycan-mediated biology, enabling translational applications in therapeutics and diagnostics.

## Methods

### Dataset

To evaluate how all-atom prediction software extrapolates to glycans, we used DIONYSUS (access date: October 8, 2024), to construct our dataset. We first selected all protein-carbohydrate complexes after the September 2021 training cutoff date used by all models. Of the 5,461 identified structures by DIONYSUS, 614 proteins were deposited in the PDB after the training date cutoff. We then clustered all 5,461 protein sequences using MMSEQS^27^ into 50% sequence identity clusters and removed any post-cutoff proteins with sequence homology with any protein published before the training date cutoff, leaving 105 structures. We then selected a single structure from each cluster, selecting the complex with the highest degree of polymerization (DP), leaving 35 protein structures. Of these 35 protein structures, 11 experimentally bind monomers, 6 experimentally bind dimers, 13 bind polymers (3+ saccharides), and 5 bind a saccharide and nucleotide triphosphate (NTP).

For each structure, we analyzed the ligand structure quality measures, notably real space R factor (RSR) and real space correlation coefficient (RSCC) (Figure S2).^23^ When these metrics weren’t available, (7TOH, 7YWF, 8CSF) we provide their root-mean-squared deviation Z-scores (RMSZ). We define the set of high-quality structures with an RSCC greater than 0.9,^23^ which contains 20 structures: 9 that bind monomers, 3 that bind dimers, 5 that bind polymers, and 3 that bind at least an NTP and a saccharide. We named our dataset the Benchmark of CArbohydrate Protein INteractions (BCAPIN).

### Prediction methodology

To provide an equivalent and biologically relevant input ligand for all structures, we generated the SMILES strings of the original PDB ligand using GlyLES^34^ (part of the Glycowork^35^ Python package). In the case of homogenous polymers, we extended the length of the original carbohydrate by a DP of 2 to provide additional biological context. AF3, Boltz-1, Chai-1 and Diffdock input a SMILES string,^7,9,12,13^, but RFAA requires an SDF file input (ligand coordinates) to perform the calculations, which we used RDKit to calculate the initial ligand coordinates. In the case of heparin binding proteins (8EDI and 7PGK), we used the SMILES retrieved from the PubChem compound instead.^36^ For the five glycosyltransferases (GTs) targets, we input both the carbohydrate(s) and NTP to the software for multi-body docking.

To replicate the process of a simple *de novo* pipeline, we ran all methods without modifications or customizations. AF3, Boltz-1, and RFAA were run with a local distribution with five random seeds using the SMILES strings (or RDKit generated SDF from the SMILES for RFAA). Chai-1 was run using the Chai-1 servers, which uses five random seeds for predictions. All confidence metrics were extracted from provided mmCIF and json files. For predicted absolute error, we rather used Boltz-1’s interface predicted distance error (ipde).

Diffdock is not an end-to-end method, therefore we ran DiffDock in two different contexts, (1) with the solved experimental structure, which we call DiffDock-*holo*, and (2) with a predicted AF3 protein structure, which we call Diffdock-*AF3*. The AF3 structure for the input into DiffDock-*AF3* was chosen as the best ranking AF3 *apo* model running from 5 random seeds. We ran both DiffDock methods using the HuggingFace server with the SMILES strings, resulting in 10 total models. On GTs with multiple ligands, we concatenate the structures of the same rank together for a singular prediction.

### Metrics

Carbohydrates differ substantially from conventional small molecules, as they range from small monosaccharides to branched polymers. We therefore selected the following metrics to analyze protein-carbohydrate complex predictions: full ligand *F*_nat_ (*F*_nat,Jull_), residue *F*_nat_ (*F*_nat,res_), ligand RMSD (LRMS), and ring-ring RMSD (rRMS).

*F*_*nat*_ is the fraction of native contacts, defined as all residue-residue contacts (any heavy atom to any heavy atom) within 5 Å:

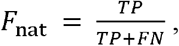

where *TP* (True Positives) is the overlap between predicted contacts and experimentally known contacts and *FN* (False negatives) are all experimental contacts not observed in the predicted structure. We use this formal definition of residue-residue contacts which we call *F*_nat,res_. In addition, as these are small molecule-like ligands, we additionally define *F*_nat,full_, which instead of carbohydrate residue-protein residue contacts, instead is the full ligand *F*_nat_, or any carbohydrate heavy atom-protein residue contacts (effectively treating the full ligand as a singular residue).

In addition to *F*_nat_, we leverage the root means squared deviation (RMSD) metric:

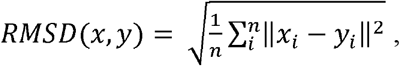

where *x*_*i*_ are the coordinates of select heavy atoms of the predicted structure and *y*_*i*_ are the same heavy atom coordinates of the experimentally determined structure after optimal superposition of the protein’s binding pocket (all residues within 10 Å of the ligand). We chose two different RMSDs to indicate the fine-grained nature of carbohydrate polymers: ligand RMSD and ring RMSD. Ligand RMSD (*LRMS*) measures the distance between the predicted and experimental structures of the ligand’s heavy atoms. For LRMS, we use the RDKit implementation that compares the maximal similar substructures. This measurement is accounts for the specific orientations (e.g. chirality and epimerization) of the carbohydrate rings.^37,38^ Ring RMSD (*rRMS*) simplifies the problem to only measuring the distance between the center of mass (COM) of each carbohydrate ring. We use a greedy implementation of *rRMS*, where each saccharide species is equivariant to any other saccharide species along the polymer chain.

We combine the four separate measurements to “DockQC,” which represents the overall quality of the predicted protein-carbohydrate structure on a scale from 0 to 1. This metric is inspired by the foundational DockQ metric for measuring protein-protein docking.^39,40^ DockQ measures on a scale from [0,1] by combining the fraction of natural contacts (*F*_*nat*_), LRMS, and interface RMS (iRMS).^39,40^

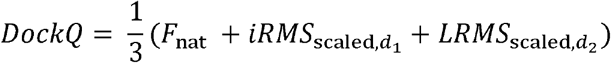

where *d*_*1*_ = 1.5 Å and *d*_*2*_ = 8.5 Å and

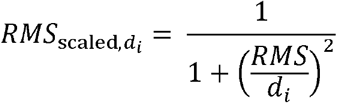

Currently, DockQ does not allow the ligands to differ in size between the crystal and predicted structure. Additionally for small molecules, DockQ only reports the LRMS value.^40^ When we reimplemented the DockQ metric with these values accounted for, we found it unrepresentative of the predictions (Table S1, Figure S1). We therefore constructed the DockQC based on the metrics as follows:

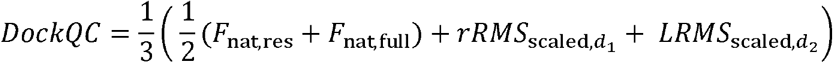

where *d*_*1*_ = 2.0 Å, *d*_*2*_ = 4.0 Å. We tuned the scaling factors of *d*_*1*_ and *d*_*2*_ to fit the DockQC into the four different categories: incorrect (DockQC < 0.25), acceptable (0.25 <= DockQC < 0.50), medium (0.50 <= DockQC < 0.80), and high (DockQC >= 0.80) (Table S1, Figure S1).

### Human proteome predictions

We selected nine proteins from the human proteome to evaluate *de novo* docking on proteomic scales, where PiCAP predicts six of these proteins as carbohydrate binding proteins and three as non-binding proteins. We used the following purported glycans for docking based on the function of each protein: GM1, Gal(β1-3)GalNAc(β1-4)[Neu5Ac(α1-3)]Gal(β1-4)Glcβ, for ganglioside binding proteins and a hybrid N-glycan for the remaining proteins, Neu5Ac(α1-6)Gal(β1-4)GlcNAc(β1-2)Man(α1-3)[Man(α1-6)[Man(α1-3)]Man(α1-6)]Man(β1-4)GlcNAc(β1-4)GlcNAcβ.

## Data Availability

The BCAPIN dataset and all model inputs, code, and analysis data are available on Github at github.com/graylab/dockqc.

## Author Contributions

S.W.C. conceived the project, performed the research, analyzed data, wrote the manuscript, and created all figures. L.L. conceived the project, performed the research, analyzed data, and wrote the manuscript. S.S.T. performed the research and analyzed data. J.J.G. conceptualized and supervised the project, analyzed data, and wrote the manuscript.

## Acknowledgements

We thank William F. Degrado for his continual support and advice on the project, identifying limitations of benchmarking when comparing experimental and computational protein-carbohydrate structures. We also thank Dr. Matthew O’Meara and Miguel Limcaoco for advice on evaluation of experimentally solved structures and of predicted structures. Computing was performed on the JHU High Performance Computing from ARCH. We thank the Rosetta Commons lab exchange travel support (LL).

