## Supplementary File 1 for "Evaluation of De Novo Deep Learning Models on the Protein-Sugar Interactome"

### DockQC parameterization

**Table S1: DockQC with different d_i_ values compared to DockQ**^1^ **on 8 targets.** Human labels for “High” quality, “Medium” quality, “Acceptable” (Acc) quality, and “Low” quality are provided.

|  | PDB | DockQ | DockQC  d_1_=3.0, d_2_=1.5 | DockQC  d_1_=5.0, d_2_=1.5 | DockQC  d_1_=4.0, d_2_=2.0 | DockQC  d_1_=3.0, d_2_=2.0 |
| --- | --- | --- | --- | --- | --- | --- |
| High | 7BLG | 0.94 | 0.93 | 0.94 | **0.94** | 0.93 |
|  | 8AD2 | 0.94 | 0.80 | 0.83 | **0.85** | 0.83 |
| Med | 7EN5 | 0.86 | 0.47 | 0.55 | **0.56** | 0.52 |
|  | 7W18 | 0.68 | 0.51 | 0.52 | **0.56** | 0.56 |
| Acc | 7F9G | 0.73 | 0.32 | 0.40 | **0.39** | 0.35 |
|  | 7EQR | 0.61 | 0.35 | 0.36 | **0.40** | 0.40 |
| Low | 8DZD | 0.37 | 0.01 | 0.02 | **0.01** | 0.01 |
|  | 8EDI | 0.61 | 0.21 | 0.23 | **0.22** | 0.21 |

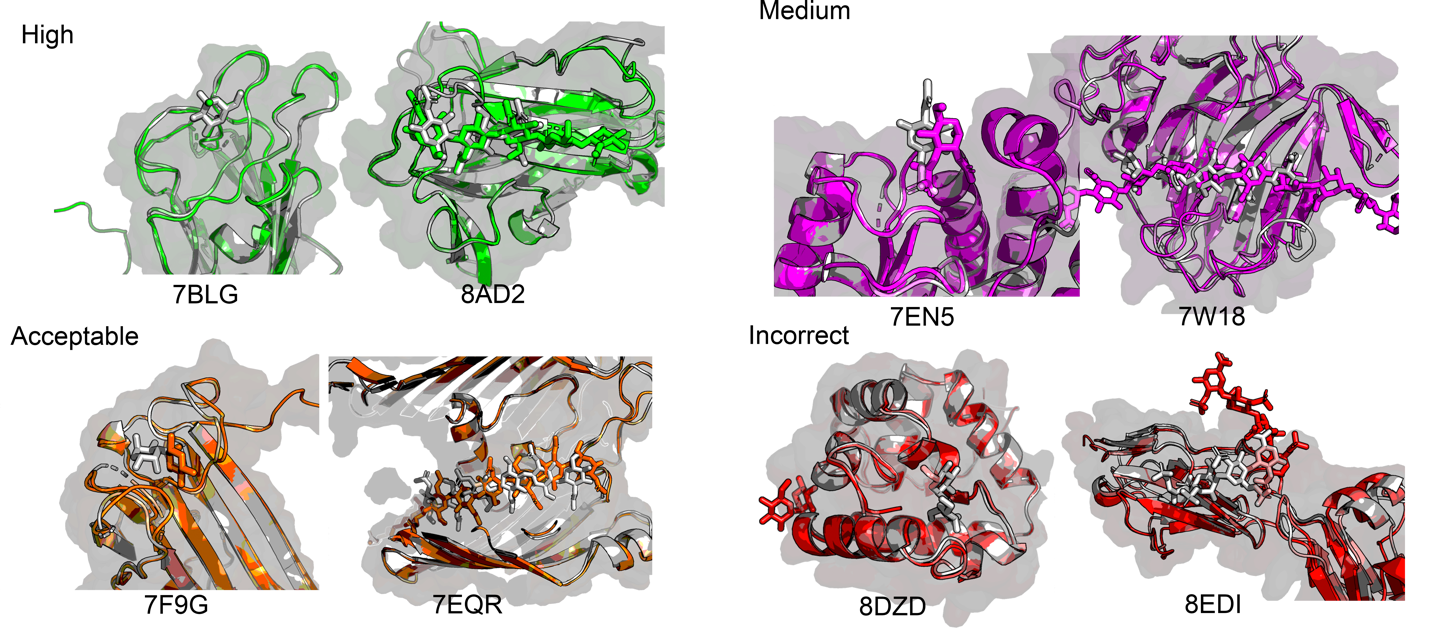

**Figure S1: Predicted protein structures deemed to be of high, medium, acceptable, and incorrect quality.**

### BCAPIN RSC and RSRR values

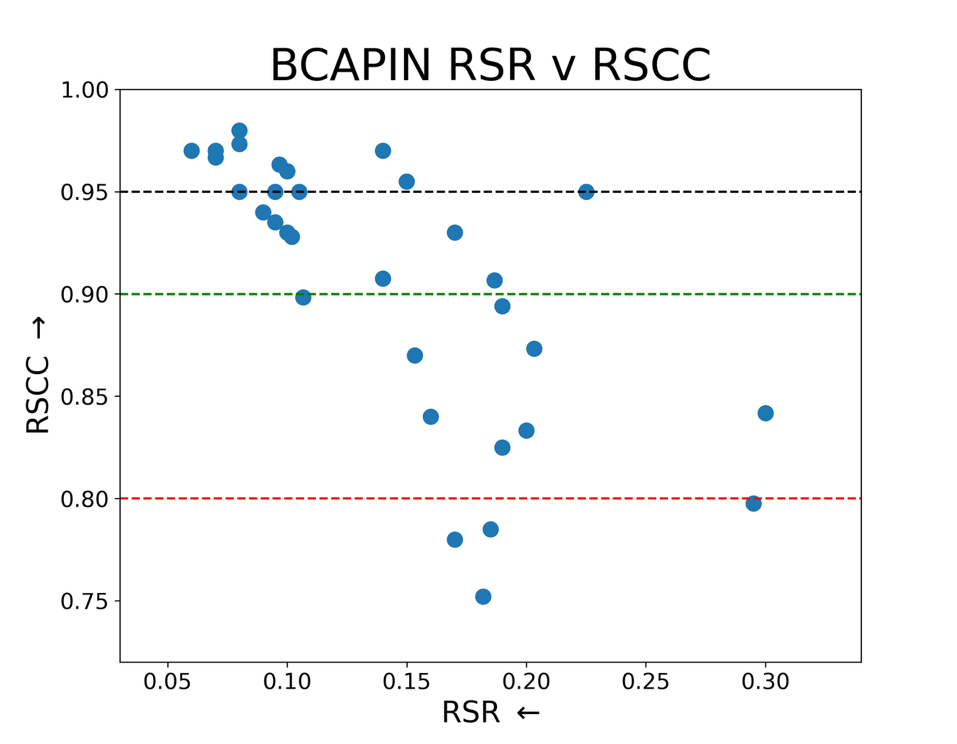

**Figure S2: Analysis of the BCAPIN real space R factor (RSR) and real space correlation coefficient (RSCC).** Lines at RSCC values of 0.95, 0.9 (the cutoff for the BCAPIN set), and 0.8 are shown.^2^

### BCAPIN confidence metrics

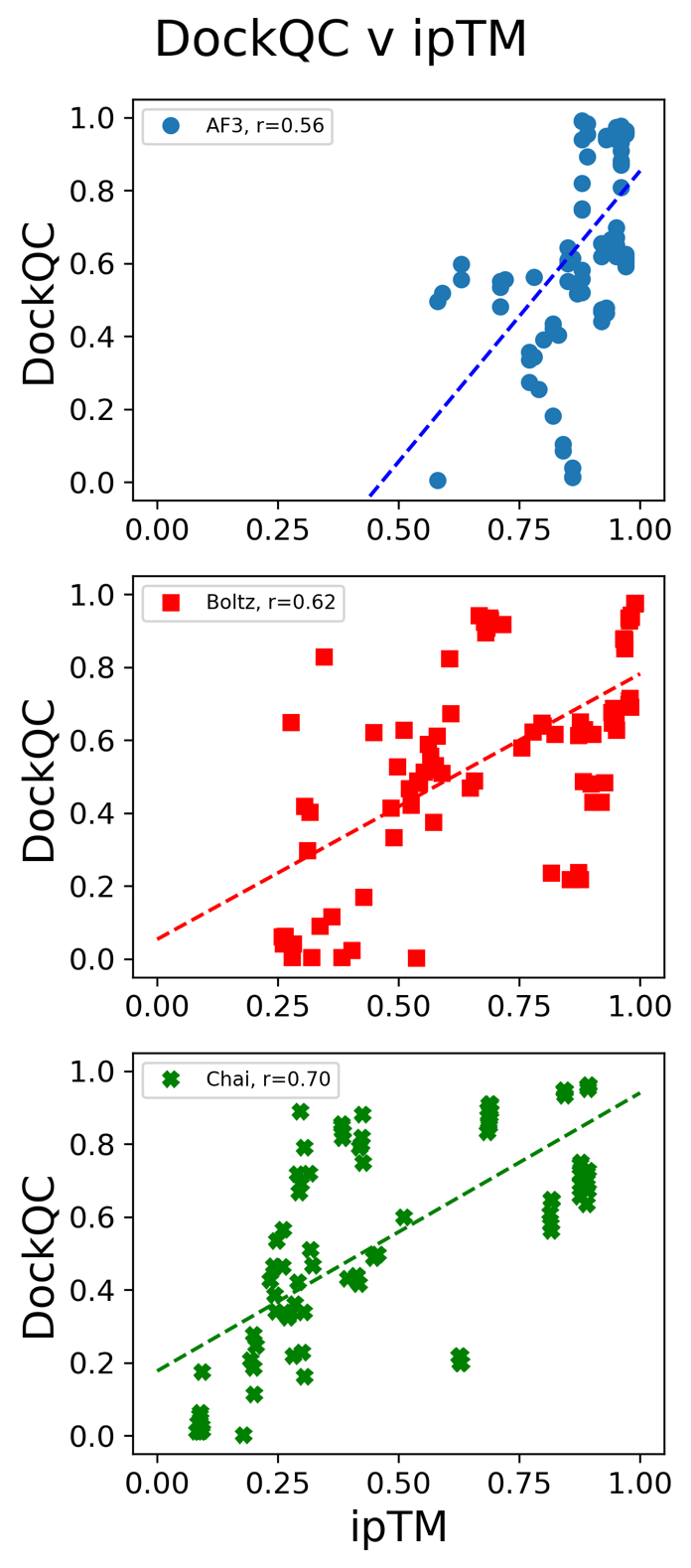

**Figure S3: Comparison of ipTM and DockQC accuracy on BCAPIN**. AF3 (blue circle), Boltz-1 (red square), and Chai-1 (green X) were the only models which reported such metric.

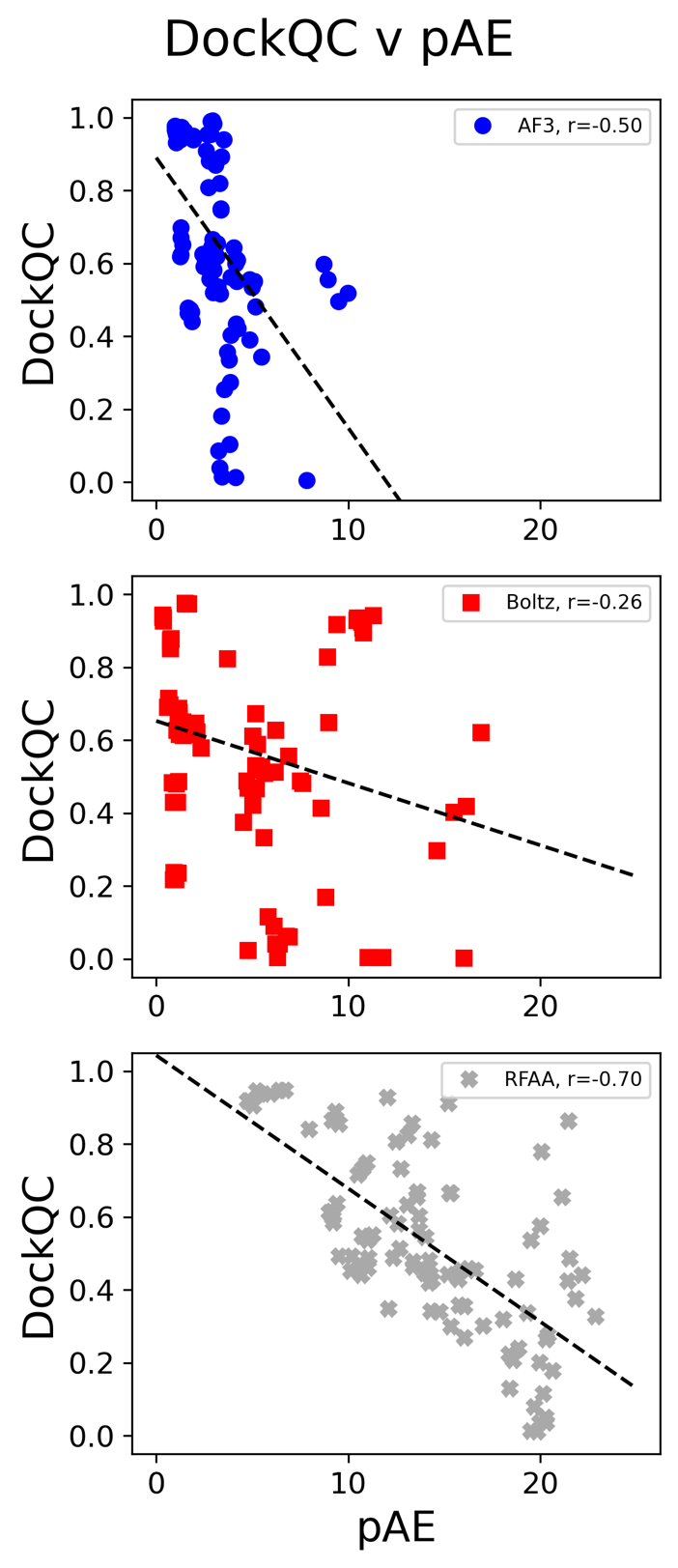

**Figure S4: Comparison of pAE and DockQC accuracy on BCAPIN**. AF3 (blue circle), Boltz-1 (red square), and RFAA (gray Diamond) were the only models which reported such metric.

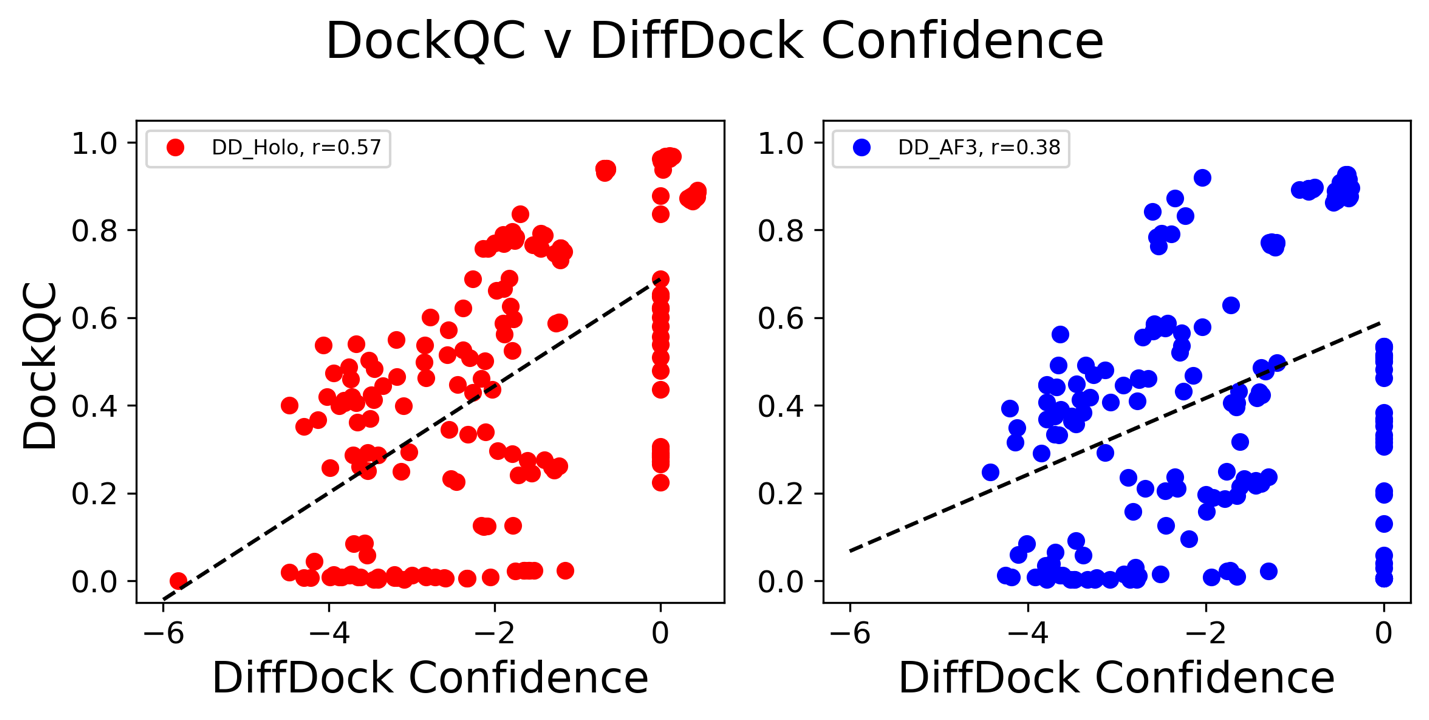
**Figure S5: Comparison of pAE and DockQC accuracy on BCAPIN**. AF3 (blue circle), Boltz-1 (red square), and RFAA (gray Diamond) were the only models which reported such metric.

### BCAPIN: Full Set analysis

To identify any discrepancies between high quality and low quality protein-carbohydrate complexes, we analyzed the total 35 structures (20 high quality, 15 low quality) below. We found similar results to analyzing only the high quality structure predictions, all models perform strongly, capturing at least 80% of all top-5 complexes with at least acceptable quality DockQC.

**Table S2: Benchmark of CArbohydrate Protein INteractions (BCAPIN) low quality proteins (RSCC < 0.9)**. The table lists the PDB 4-letter ID, protein name, UniProt ID, glycan input string for GlyLES, and any secondary ligands if present.

| PDB | Protein Name | UniProt | | GlyLES Input String | | Secondary Ligand |
| --- | --- | --- | --- | --- | --- | --- |
| 7eqr | Chitoporin | L0RVU0 | | GlcNAc(b1-4)GlcNAc(b1-4)GlcNAc(b1-4)GlcNAc(b1-4)GlcNAc(b1-4)GlcNAc(b1-4)GlcNAc(b1-4)GlcNAc(b1-4)GlcNAc | |  |
| 7lvy | UDP-glycosyltransferase 203A2 | T1K1R5 | | Glc | | UDP |
| 7p8g | Glucosyl-3-phosphoglycerate synthase | K5B7Z4 | | Glc | |  |
| 7pgk | Hedgehog-interacting protein | Q96QV1 | | Heparin_analog: PDB SMILES used | |  |
| 7pug | GH115 | N/A | Xyl(b1-4)Xyl(b1-4)Xyl(b1-4)Xyl(b1-4)Xyl(b1-4)Xyl(b1-4)Xylb | | |  |
| 7toh | SGNH hydrolase | A0A5M4AV20 | | GlcA4Me(a1-2)Xylb | |  |
| 7tvp | Ciral AMG chitosanase | Unk | GlcNAc(b1-4)GlcNAc(b1-4)GlcNAc(b1-4)GlcNAc(b1-4)GlcNAc | | |  |
| 7vu1 | Chitoporin | P75733 | | GlcNAc(b1-4)GlcNAc(b1-4)GlcNAc(b1-4)GlcNAc(b1-4)GlcNAc(b1-4)GlcNAc(b1-4)GlcNAc | |  |
| 7vwb | 17 kDa phloem lectin | Q8LK69 | | Gal(b1-4)GlcNAca | |  |
| 7xtn | α-1,3-mannosyl-glycoprotein 4-β-N-acetylglucosaminyltransferase A-like isoform X1 | A0A6J2K041 | | GlcNAc | |  |
| 7ywf | Dirigent protein | Q306J3 | | Gal(a1-3)Galb | |  |
| 8ad2 | Nictaba | Q94EW1 | | GlcNAc(b1-4)GlcNAc(b1-4)GlcNAc(b1-4)GlcNAc(b1-4)GlcNAc | |  |
| 8csf | WbbB D232C-Kdo adduct | Q6U8B0 | | | Rha(a1-3)GlcNAcb | Ligand 2: GDP  Ligand 3: Kdo |
| 8edi | Netrin receptor unc-5 | Q26261 | | | Heparin_analog: PDB SMILES used |  |
| 8ped | Alginate lyase | [A0A7I9C8Z1](https://www.uniprot.org/uniprot/A0A7I9C8Z1) | ManA(b1-4)ManA(b1-4)ManA(b1-4)ManA(b1-4)ManA(b1-4)ManA(b1-4)ManA(b1-4)ManA(b1-4)ManA(b1-4)ManAb | | |  |

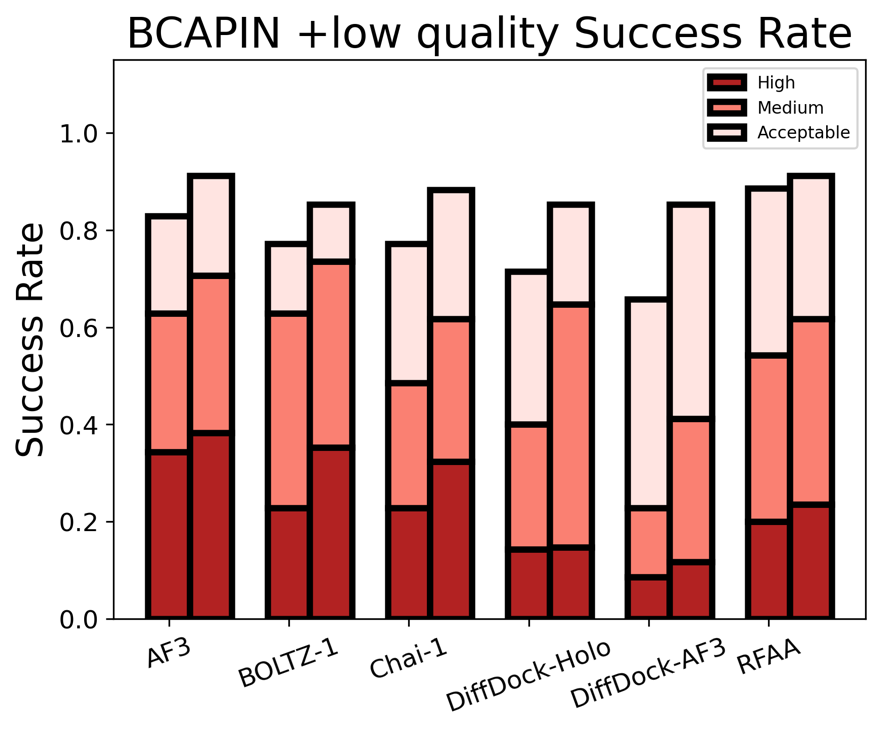

**Figure S6: Analysis of the BCAPIN test set, including the low quality structures (RSCC < 0.9) (n=35).** Top-1 (left) and top-5 (right) predictions of each method.

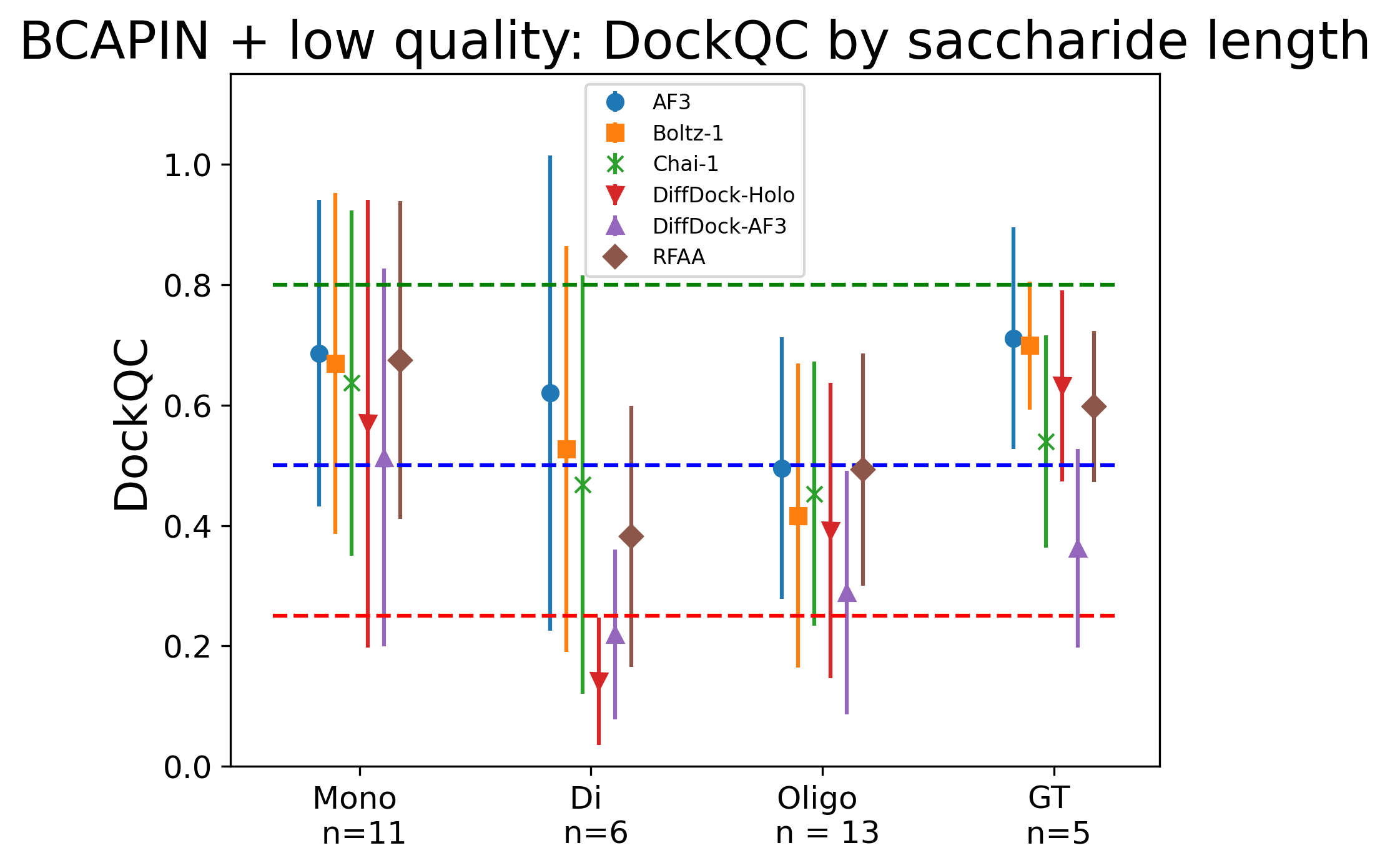

**Figure S7: Comparison of average and standard deviation DockQC of predicted structures versus saccharide length for BCAPIN set with the low quality structures (RSCC < 0.9).** We group saccharide length into a degree of polymerization (DP) of 1 (mono), 2 (di), and 3+ (oligo), and further group all glycosyltransferases (GTs) together that require multiple inputs (e.g. a saccharide and NTP) and with the number of proteins in each group listed. We also show the DockQC cutoffs between acceptable (red), medium (blue), and high (green) quality structures. Top-1 prediction on the gold set with AF3 (blue circle), Boltz-1 (orange square), Chai-1 (Green X), Diffdock-*holo* (red triangle), Diffdock-*AF3* (purple triangle), and RFAA (brown diamond).
